# Genomic regulation of chemo-mechanical stability in plant-derived extracellular vesicles: a multiscale model of composite reinforcement

**DOI:** 10.64898/2026.07.01.735926

**Authors:** Jagir R. Hussan, Anand Rampadarath, David P. Nickerson, Peter J. Hunter

## Abstract

Plant-derived extracellular vesicles (PDEVs) have emerged as superior candidates for oral drug delivery, exhibiting a gastrointestinal survivability that significantly exceeds that of mammalian exosomes or synthetic liposomes. However, the biophysical rules governing how plant genomic regulation translates into this exceptional mechanical resilience remain unknown. Here, we present a predictive multiscale model of plant-derived extracellular vesicles, linking a parameterised genetic state space to emergent mesoscale mechanics via supra-molecular coarse-grained molecular dynamics (SCG-MD). We demonstrate that the upregulation of sterol methyltransferases (SMT) during the plant’s theoretical Defence state drives the formation of a phase-separated composite architecture, where rigid domains occupying approximately 36% of the membrane surface area effectively arrest crack propagation. This state achieves a critical rupture tension of 367.0 ± 0.7 mN m^−1^ corresponding to a 39% increase over the wild-type Ripening state. Crucially, we find that chemical composition alone is insufficient for this reinforcement; vesicles with actively sorted lipid domains (Seeded topology) outperform randomised mixtures (Spontaneous topology) by 23% at identical concentrations. Furthermore, while fluid vesicles stiffen reactively under gastric acid shock (pH 2.5) due to the steric jamming of thermodynamically neutralised headgroups, the Defence state exhibits mechanical homeostasis. These findings suggest that PDEVs function as genetically tunable composite materials, offering a design blueprint for next-generation bio-inspired drug delivery vectors. Ultimately, these theoretical indices provide a predictive biophysical framework awaiting empirical confirmation via *in vitro* nanomechanical assays.

## 1 Introduction

The development of efficient oral drug delivery vectors remains a formidable challenge in nanomedicine. To succeed, a carrier must navigate the human gastrointestinal (GI) tract, a harsh sequence of physicochemical barriers characterised by extreme pH gradients (pH 1.5–7.5), bile salts, and enzymatic degradation. ^1,2^ While synthetic liposomes and mammalian exosomes frequently destabilise or fuse prematurely under these conditions, plant-derived extracellular vesicles (PDEVs) have demonstrated a remarkable intrinsic ability to survive gastric transit and deliver bioactive cargo across kingdom boundaries. ^3,4^ This durability identifies PDEVs not merely as biological messengers, but as naturally evolved armoured containers that outperform current synthetic standards.

This mechanical resilience is likely a consequence of the unique evolutionary pressures exerted on plant cells. Unlike mammalian cells protected by homeostatic tissues, plant cells are directly exposed to abiotic stress and substantial osmotic fluctuations. Consequently, they have evolved potent genomic mechanisms to remodel their lipid membranes in response to environmental stimuli. During fruit ripening, master regulators such as NAC and MADS-box transcription factors coordinate a physiological shift toward fluidity, upregulating fatty acid desaturases (FADs) to facilitate the secretion of aroma volatiles. ^5,6^ Conversely, in response to pathogen attack or physical injury, plants trigger a *Defence state*, upregulating enzymes like sterol methyltransferases (SMTs) to rigidify the cell membrane and form barrier structures. ^7–9^ Because PDEVs originate either from the outward budding of the plasma membrane (ectosomes) or from the fusion of the multivesicular body with the plasma membrane (exosomes), they inevitably inherit these stress-tuned lipid profiles ^10^.

However, a critical gap remains in our understanding of this process. While the molecular biology of plant stress and the chemistry of lipid bilayers are individually well-documented, the *biophysical principles* connecting the two are missing. We do not currently know how a specific shift in gene expression (e.g., SMT up-regulation) quantitatively translates into the continuum mechanical properties of the secreted vesicle. Furthermore, it is unclear whether the stability of PDEVs arises simply from their bulk chemical composition (a stiffer ingredients hypothesis) or if it requires a specific supramolecular organisation, such as the formation of load-bearing nanodomains. In soft-matter physics, composite reinforcement, whereby discontinuous rigid inclusions impede crack propagation within a compliant matrix, is a well-established toughening mechanism ^11^. We hypothesise that PDEVs exploit an analogous strategy, utilising phase-separated domains as a dispersed load-bearing architecture, similar to the high lysis tensions observed in engineered raft-forming mixtures. ^12,13^

In this work, we address this gap by developing a multiscale chemo-mechanical model that connects the highly conserved genomic regulation of plant stress responses to the physical stability of secreted PDEVs. We parameterise the genetic expression space to map theoretical Ripening and Defence profiles directly to a coarse-grained molecular dynamics framework. By subjecting these digital vesicles to simulated osmotic swelling and gastric acid shock (pH 2.5), we isolate the electrostatic and mechanical stressors of the GI tract to test the hypothesis that PDEV stability is an emergent property of composite reinforcement. While native gastrointestinal transit involves a complex interplay of bile salts and enzymatic degradation, our model explicitly evaluates osmotic- and pH-driven survivability to determine baseline membrane mechanics.

We identify a mechanical hierarchy where the Defence phenotype significantly outperforms the wild-type baseline. Crucially, our results demonstrate that composition is not enough; the high rupture tensions observed in nature can only be replicated if lipids are actively sorted into phase-separated domains, rather than randomly mixed. This implies that the plant cell utilises a tunable armour strategy, actively engineering the topology of its vesicles to withstand environmental stress. These findings provide a theoretical basis for the exceptional stability of plant vesicles and offer a biomimetic blueprint for engineering mechanically homeostatic drug delivery systems.

## 2 Methods

### 2.1 Parameterisation of genetic states

To construct a computationally tractable state space, we define the vesicle’s chemo-mechanical profile as a function of three rate-limiting enzymes: fatty acid desaturase (*G*_*FAD*_), sterol methyl-transferase (*G*_*SMT*_), and phospholipase D (*G*_*PLD*_). These regulators were selected because they represent the primary biophysical levers plants utilise to remodel membrane mechanics under stress ^14–16^. Specifically, *G*_*FAD*_ dictates acyl chain polyunsaturation, governing core fluidity and bending compliance; *G*_*SMT*_ regulates phytosterol synthesis, driving thermodynamic condensation and in-plane stiffening; and *G*_*PLD*_ mediates the production of anionic phosphatidic acid, tuning headgroup electrostatics. While these lipid metabolic pathways are intimately coupled *in vivo*, we adopt a simplifying modelling assumption treating them as mathematically orthogonal variables. This abstraction establishes a theoretical phase space, allowing us to continuously map the complex biological stress response and explicitly isolate the distinct mechanical contribution of each pathway:

1. **State A (Ripening):** Defined by high *G*_*FAD*_ activity, representing the wild-type maturation profile. This upregulates polyunsaturated fatty acid production (*S*_*unsat*_ ≈ 80%) via FAD activity ^6^. At the mesoscale, polyunsaturation fundamentally alters the mechanical properties of the lipid bilayer, creating a hyper-fluid matrix with reduced bending rigidity ^12,17^. Biophysically, this compliant state lowers the energy barrier for fusion and lysis, theoretically facilitating the secretion of cell-wall degrading enzymes.
2. **State B (Defence):** Defined by setting *G*_*SMT*_ to the upper boundary of our simulated parameter space (4.0-fold relative expression). We establish this theoretical maximum as a modelling assumption to represent an extreme stress-induced phenotype. Biologically, this state mimics the stress-induced accumulation of phytosterols and the concurrent exclusion of phospholipids observed in plant membrane micro-domains ^18^. Biophysically, this sustained sterol enrichment is required to surpass the critical concentration threshold necessary to trigger lateral phase separation and the macroscopic percolation of liquid-ordered (*L*_*o*_) domains, a phenomenon well-documented in model biomembranes ^19^. Simultaneously, *G*_*PLD*_ activity modulates the concentration of anionic phosphatidic acid, influencing electrostatic stability. To simulate the formation of functional plant nanodomains ^20^, the cohesive interaction energy driving *L*_*o*_ phase demixing is parameterised to *ε*_*Lo*_ = 15.0 kJ mol^−1^ (≈6*k*_*B*_*T*). This specific value operates as a necessary modelling optimization: it provides sufficient thermodynamic driving force to overcome entropic mixing, while remaining below the threshold that would induce artifactual crystalline freezing, thereby ensuring realistic mesoscale membrane fluidity. ^13^

### 2.2 Chemo-mechanical mapping

To translate gene expression into lipid bilayer mechanics, we utilise a linear constitutive mapping parameterised using literature values from micropipette aspiration ^12^ and vesicle fluctuation analysis ^21^. We selected 1-palmitoyl-2-oleoyl-sn-glycero-3-phosphocholine (POPC) as the computational proxy for the fluid lipid matrix. While the native plant plasma membrane possesses a highly complex lipidome ^22,23^, POPC serves as an ideal thermodynamic baseline for the bulk liquid-disordered (*L*_*d*_) phase. Its asymmetric saturation (16:0/18:1) ensures it remains fluid at physiological temperatures while reliably supporting sterol-driven liquid-ordered (*L*_*o*_) phase separation. This allows us to anchor our elastic network to rigorously validated experimental mechanical moduli (*K*_*base*_ ≈ 240 mN m_−1_).

As a first-order heuristic approximation, we model the effective mechanical network modulus of the vesicle (*K*_*e f f*_) as a linear combination of these competing transcriptomic drivers:

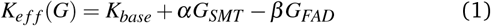

where *K*_*base*_ = 240 mN m^−1^ represents the baseline modulus of a fluid POPC bilayer. The genetic parameters *G*_*SMT*_ and *G*_*FAD*_ represent continuous relative fold-expression levels (0.5 to 4.0). The stiffening coefficient *α* = 120 mN m^−1^ approximates the condensing effect of sterols, which increase membrane moduli 2-to-3-fold ^24^. Due to a lack of macroscopic mechanical data for plant-specific sterol mixtures, we apply this cholesterol-derived coefficient as a computational proxy. This simplifying assumption is justified by comparative studies demonstrating that phytosterols (e.g., *β*-sitosterol) exert functionally analogous condensing effects on fluid matrices ^23^.

Conversely, while true thermodynamic area compressibility (*K*_*A*_) is largely invariant across differing levels of acyl chain polyunsaturation ^25^, polyunsaturation dramatically reduces bending rigidity and macroscopic lysis tension. We use a 2D SCG-MD mesh which lumps in-plane elasticity and out-of-plane compliance into a single harmonic bond network (*k*_*bond*_ ≈ 0.5*K*_*e f f*_), to mitigate this we introduce the softening coefficient *β* = 60 mN m^−1^. This acts as a phenomenological penalty to the global network modulus, accurately capturing the fluidised, rupture-prone phenotype driven by *G*_*FAD*_ upregulation without violating the physical invariance of intrinsic *K*_*A*_. Note that, *G*_*PLD*_ acts strictly on the electrostatic potentials (*U*_*dh*_) rather than the mechanical elastic network and hence does not appear in Equation (1).

### 2.3 Supra-molecular coarse-grained MD

We implemented a supra-molecular coarse-grained (SCG) model using the OpenMM framework. ^26^ The vesicle is modelled as a spherical shell of radius *R* = 20 nm. To ensure thermodynamic stability and prevent steric overlap artefacts, particle density was calibrated to *ρ* = 0.5 nm^−2^ (*N* ≈ 2500 particles).

#### 2.3.1 Simulated osmotic inflation

To quantify mechanical stability, we subjected the vesicles to simulated osmotic inflation. This method explicitly mimics the turgor pressure or osmotic differential experienced by a vesicle in a hypotonic environment. We applied a radial force, 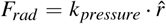, to all particles. To maintain a constant kinetic environment, *k*_*pressure*_ was incrementally ramped at a rigorous loading rate of 0.25 kJ mol^−1^nm^−1^ps^−1^ (incremented every 500 integration steps at a 4 fs timestep). This generates a uniform surface tension *γ* equivalent to a transmembrane pressure difference. Failure is defined as the macroscopic yield point, the stress at which the vesicle undergoes irreversible plastic expansion (> 10% radial strain). At the supra-molecular scale, this corresponds to the yielding of the harmonic bond network and subsequent nanopore nucleation ^27^. The critical rupture tension is calculated via the Young-Laplace law: *γ*_*crit*_ = *P*_*yield*_ *R/*2. It should be noted that because this simulated radial force is applied uniformly and deterministically across a harmonic bond network, the variance in rupture tension across replicates is minimum. In this coarse-grained regime, failure is dominated by the deterministic elastic limit of the specified harmonic springs rather than stochastic, thermally activated pore nucleation.

**Figure 1.**
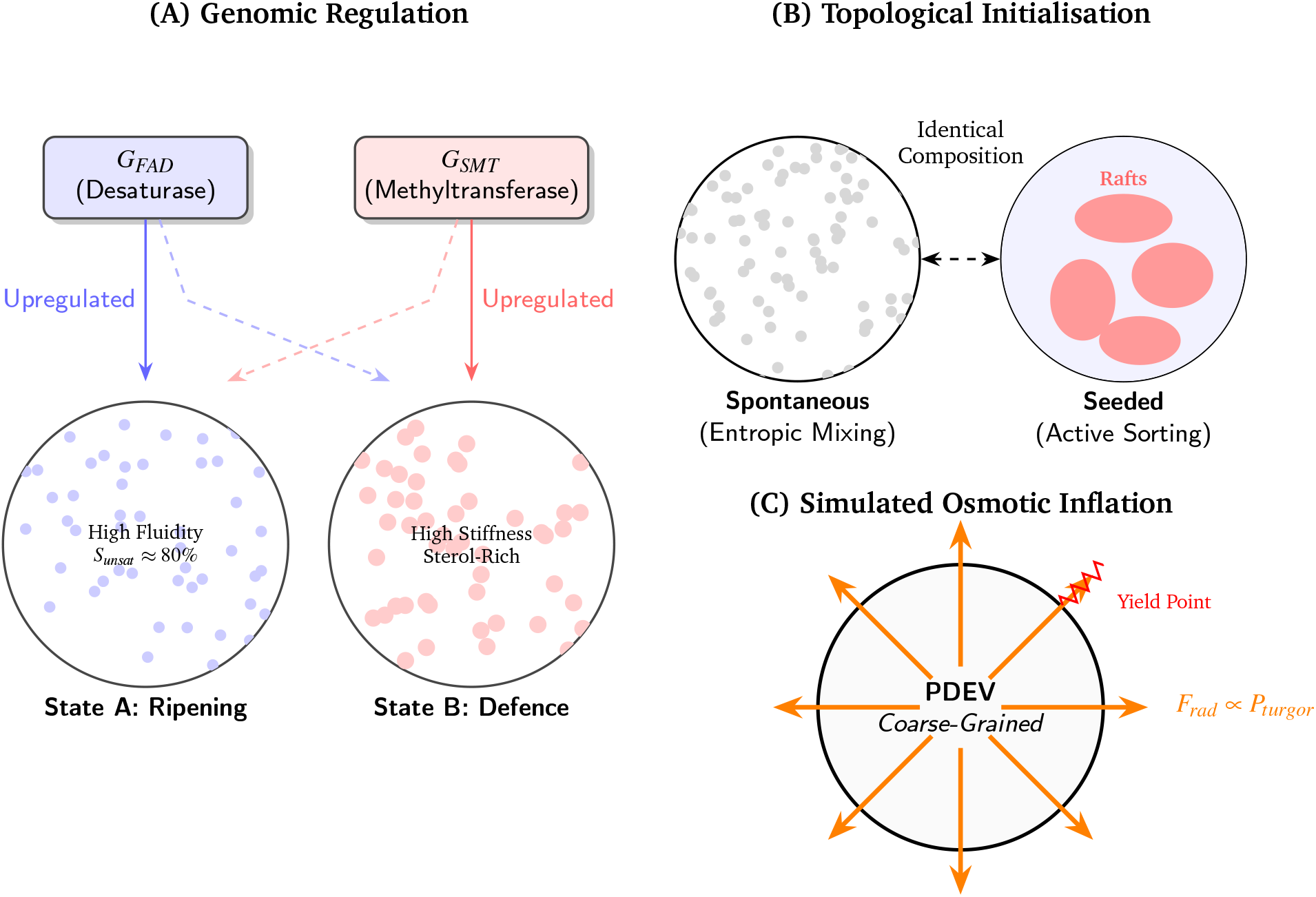
Multiscale modelling framework. **(A) Genomic regulation:** Transcription factors drive the transition from a fluid Ripening state (high FAD activity, polyunsaturated lipids) to a rigid Defence state (high SMT activity, sterol accumulation). **(B) Topological initialisation:** To test the role of organisation, identical lipid mixtures are instantiated either as Spontaneous (random entropic mixing) or Seeded (active sorting into pre-nucleated raft domains). **(C) Simulated osmotic inflation:** Vesicles are subjected to a radial force (*F*_*rad*_) mimicking turgor pressure to quantify the critical yield point and rupture tension (*γ*_*crit*_).

#### 2.3.2 Topological initialisation: sorting vs. mixing

A key objective of this study is to determine if mechanical reinforcement requires specific lipid organisation. We therefore defined two initialisation modes:

1. **Seeded (Active sorting):** This mode simulates the active biological sorting of lipids (e.g., in the Golgi or via plasma membrane flippases) into pre-nucleated domains. We initialised the topology using a cluster-growth algorithm to create phase-separated armoured domains, accelerating the formation of liquid-ordered (*L*^*o*^) structures.
2. **Spontaneous (Entropic mixing):** This mode acts as a control, representing a vesicle where lipids are randomly distributed and allowed to self-assemble solely via entropic mixing, without active cellular organisation.

#### 2.3.3 Force fields

The potential energy landscape includes:

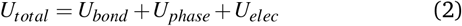

where *U*^*bond*^ represents the elastic network scaled to the local modulus (*k*^*bond*^ = 0.5*K*^*A*^), and *U*^*phase*^ is a Lennard-Jones potential with a well depth *ε*^*Lo*^ = 15.0 kJ mol^−1^ favouring sterol-sterol association. ^13^ To isolate the mechanical consequences of lipid protonation from complex solvent electrokinetics, pH-dependent electrostatics were treated using a static partial-charge approximation rather than dynamic charge regulation. Prior to topology initialisation, the net anionic charge of phosphatidic acid (*q*_*anionic*_) was scaled according to the bulk pH using the Henderson-Hasselbalch equation (*pK*_*a*_ ≈ 4.0). Consequently, at pH 2.5, anionic headgroups are modelled as thermodynamically neutral (*q* ≈ −0.03). Electrostatic repulsion was computed via a screened Coulomb potential with the Debye length (*λ*_*D*_ = 0.8 nm) fixed as a global static parameter. This decoupling ensures that the Debye length strictly represents the background ≈150 mM ionic strength of gastric fluid, while explicitly excluding non-linear Poisson-Boltzmann effects or dynamic counter-ion condensation during membrane deformation. Therefore, the stress-stiffening observed in fluid vesicles under acid shock is driven purely by the sudden loss of inter-lipid electrostatic repulsion and subsequent steric jamming of the neutralised continuum.

## 3 Results

### 3.1 Mechanical hierarchy of states

Simulated osmotic inflation experiments confirmed that the genetic parameterisation translates into distinct mechanical phenotypes. The wild-type **Ripening State (A)** exhibited a critical rupture tension of *γ*_*crit*_ = 264.1± 0.5 mN m^−1^ (SEM). In contrast, the theoretical **Defence State (B)** achieved *γ*_*crit*_ = 367.0 ± 0.7 mN m^−1^. This represents a 39% increase in rupture tension (*p* ≪ 0.001, Cohen’s *d* = 28.1). This result validates that upregulation of the sterol pathway (*G*_*SMT*_) is sufficient to push the vesicle into a hyper-stable regime, capable of withstanding osmotic pressures significantly higher than the ripening baseline.

**Figure 2.**
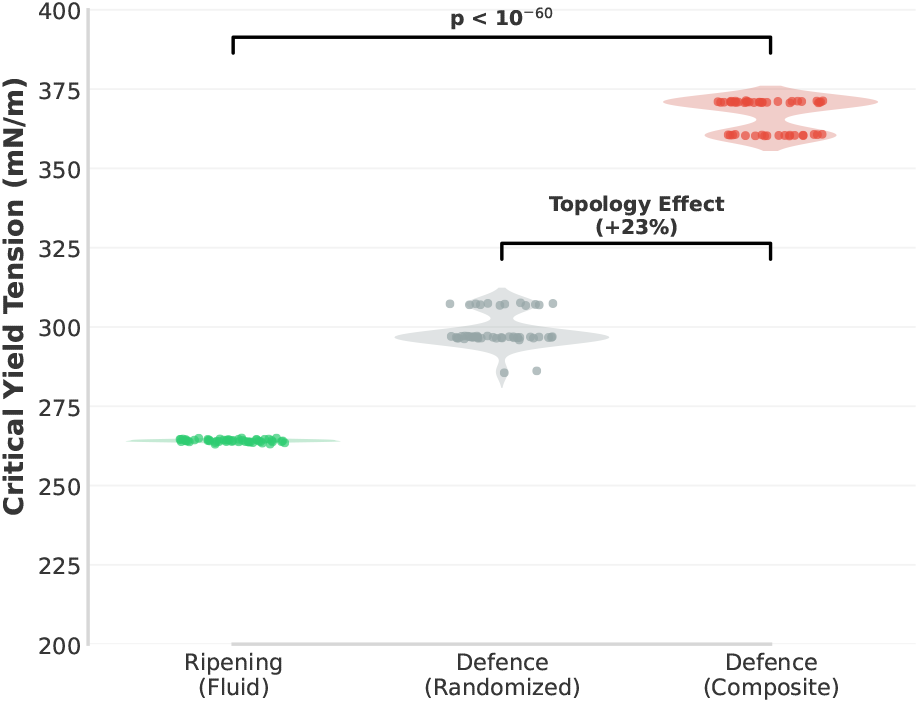
Mechanical hierarchy of PDEV states (N=50). The Defence state (`Seeded`) significantly outperforms both the Ripening state and the randomised control, demonstrating topological reinforcement.

#### Role of lipid organization

To dissect the contribution of topology, we compared vesicles where lipids were actively sorted into domains (`Seeded`) against those formed by random entropic mixing (`Spontaneous`). While the Defence composition provided a baseline strength advantage in both cases, the **Seeded** phenotype significantly outperformed the **Spontaneous** control (367.0 mN m^−1^ vs. 298.8 mN m^−1^; *p* ≪ 0.001). This 23% performance gap indicates that chemical composition alone is insufficient to explain the full stability of PDEVs; the specific arrangement of lipids into phase-separated domains is a critical mechanical factor.

### 3.2 Topological reinforcement mechanism

To understand the physical origin of this reinforcement, we analysed the connectivity of the sterol-rich phase using percolation theory. The **Seeded** Defence state exhibited a mean percolation score of *S*_*max*_ = 36.1% ± 8.3%. Visually and topologically, this corresponds to a composite architecture: discontinuous rigid plates embedded in a fluid matrix.

In contrast, the **Spontaneous** control vesicles converged to a monolithic percolation regime (*S*_*max*_ ≈ 49.6%), representing a single, continuous rigid network. Despite containing the exact same lipids, this monolithic structure was 18% weaker than the composite architecture. This suggests that PDEVs function as composite materials: the fluid matrix dissipates energy, while the discontinuous rigid domains act as armour plates that arrest crack propagation. When the rigid phase becomes continuous (monolithic), cracks can propagate unimpeded through the brittle network, leading to premature failure.

**Figure 3.**
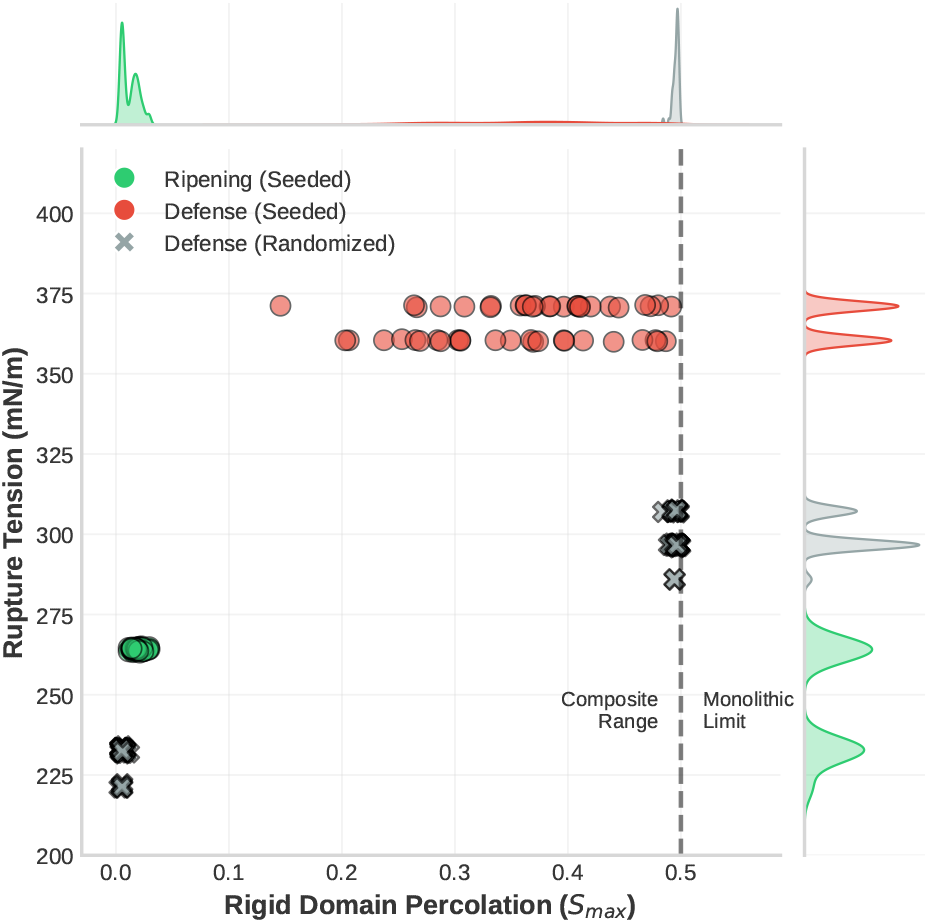
Figure 3 Percolation analysis reveals distinct topological regimes. *S*_*max*_ is defined as the fraction of total membrane particles belonging to the largest connected rigid component. A transition from *S*_*max*_ ≈ 0.15 (Dispersed) to *S*_*max*_ ≥ 0.45 (Monolithic limit for a 50% rigid composition) indicates the formation of a global percolating network. The Defence state (Seeded) clusters at *S*_*max*_ ≈ 36%, indicating discontinuous armoured domains, whereas the Randomised control (Spontaneous) clusters at *S*_*max*_ ≈ 50%, indicating a monolithic percolating network.

### 3.3 Acid shock and mechanical homeostasis

Subjecting the vesicles to simulated gastric fluid (pH 2.5) revealed a divergent response grounded in electrostatics. The fluid Ripening State (A) exhibited a stress-stiffening response, with tension increasing by 8.9% due to the protonation of anionic lipids and subsequent electrostatic jamming.

Conversely, the Defence State (B) exhibited **mechanical homeostasis**, with tension shifting by only 2.4% under identical acid shock. The sterol-rich composite network provides a stabilising mechanical scaffold that buffers the membrane against these electrostatic fluctuations, maintaining structural integrity across the GI pH gradient.

## 4 Discussion

The central question motivating this study was to identify the biophysical principles that allow plant-derived extracellular vesicles (PDEVs) to survive gastrointestinal transit, an elusive feature among synthetic liposomes. Our results suggest that this resilience is not merely a consequence of stiffer ingredients but is an emergent property of a tunable composite topology. By simulating the genomic transition from a fluid Ripening state to a sterol-rich Defence state, we found that the plant cell can increase vesicle rupture tension by 39% without completely sacrificing fluidity. This confirms our initial hypothesis that the upregulation of sterol methyltransferases (*G*_*SMT*_) acts as a master mechanical switch, converting the membrane from a secretory vehicle into an armoured stress-response unit.

However, the most significant insight arises from the discrepancy between our `Seeded` and `Spontaneous` models. A critical determination was whether the stability of PDEVs arises merely from bulk chemistry or necessitates a specific supramolecular organisation. The fact that randomised vesicles were 23% weaker than their phase-separated counterparts, despite having identical lipid compositions, provides a definitive answer: topology matters. In soft matter, a monolithic rigid network is often brittle. Our topological analysis (*S*_*max*_) reveals that when a rigid phase is allowed to spontaneously percolate and occupy ~ 50% of the total vesicle surface area, it forms a continuous network where local ruptures rapidly propagate as catastrophic pores. In contrast, the Seeded Defence state forms a discontinuous composite covering only ~ 36% of the global surface area. This architecture effectively arrests nanopore expansion; the rigid domains act as steric barriers that isolate localised lipid lattice yield events, preventing catastrophic macroscopic failure and allowing the vesicle to withstand higher critical strains ^12^.

**Figure 4.**
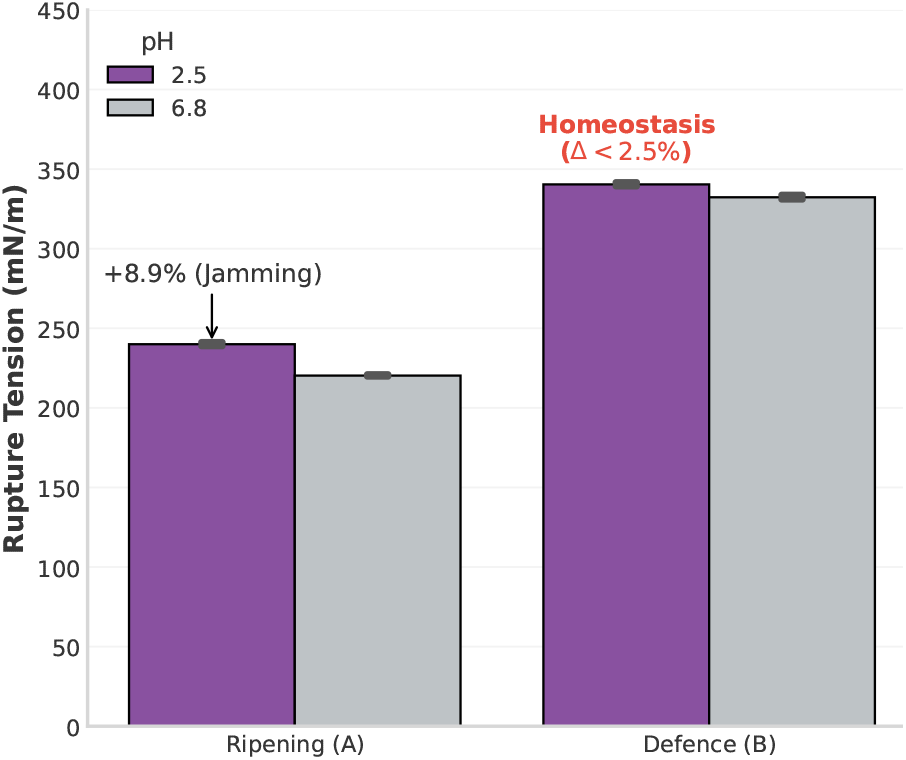
Environmental stability under acid shock. The Defence state maintains mechanical homeostasis (Δ < 2.5%), while the Ripening state reacts significantly (Δ ≈ 9%) to pH changes.

**Figure 5.**
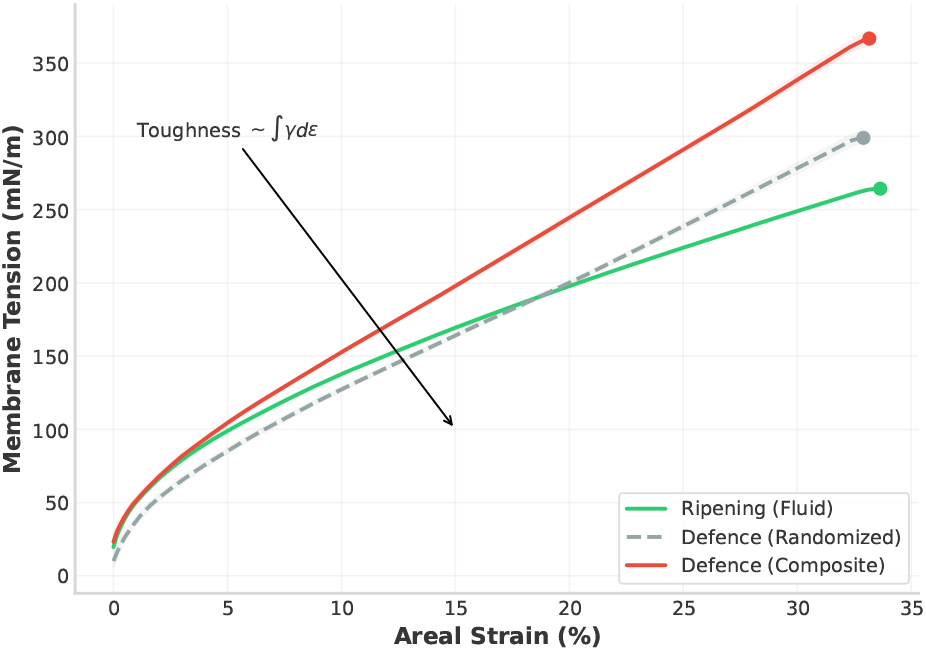
Comparative stress-strain trajectories. State B (Defence, Composite) exhibits a higher yield point and greater area under the curve (Toughness) compared to both the fluid State A and the randomised control.

This topological requirement carries significant biological implications. It suggests that the exceptional stability of PDEVs is not a passive outcome of lipid biosynthesis but likely requires active regulation during biogenesis. For a plant cell to produce these armoured vesicles, it cannot simply dump sterols and phospholipids into the cytoplasm and rely on entropic mixing. Instead, our model implies the existence of active sorting mechanisms, potentially driven by flippases or curvature-sensing proteins in the Golgi apparatus or plasma membrane which could pre-nucleate these domains before vesiculation. This aligns with recent observations of functional nanodomains in plant plasma membranes ^20^ and suggests that the plant cell effectively manufactures composite materials to survive abiotic stress.

The functional advantage of this composite architecture is further highlighted by its response to environmental shock. While the fluid Ripening state reacted to low pH (simulated gastric acid) by stiffening, the Defence state exhibited mechanical homeostasis, maintaining a constant tension profile regardless of pH. Mechanistically, this stiffening in the Ripening state is driven by electrostatic jamming. By utilising a static partial-charge approximation and a fixed Debye screening length, our model explicitly isolates the thermodynamic neutralization of the lipid headgroups from complex solvent electrokinetics. As anionic phosphatidic acid (pKa ≈ 4.0) ^28^ is initialised at pH 2.5, its assigned net negative charge approaches zero. This demonstrates that the observed reactive stiffening is fundamentally a steric phenomenon. Consequently, while the linear Debye-Hückel approximation typically breaks down in highly concentrated acidic environments due to non-linear Poisson-Boltzmann effects ^29^, the primary physical driver in our model is the outright neutralisation of the lipid head-groups. Because the screened repulsive energy scales proportionally with the product of the discrete charges (*U*_*dh*_ ∝ *q*_*i*_*q* _*j*_), the energetic term mathematically vanishes as the charge approaches zero. In this limit of neutral surface charge, complex Poisson-Boltzmann non-linearities become physically irrelevant. This abrupt loss of charge eliminates inter-lipid electrostatic repulsion, allowing a sudden decrease in the effective area per lipid and increasing local packing density, which forces the fluid continuum into a jammed, dynamically arrested state ^30^. For a drug delivery vector, this predictability is crucial. A carrier that stiffens reactively may become brittle or alter its release kinetics unpredictably in the stomach. The mechanical homeostasis observed in the Defence phenotype suggests that nature has evolved a structure capable of buffering the internal cargo against the extreme physicochemical fluctuations of the external environment.

It is important to acknowledge the limitations of this predictive model. Firstly, to maintain computational tractability within the supra-molecular coarse-grained (SCG) framework, our simulations utilise a vesicle radius of 20 nm (*N* ≈ 2500 particles). Crucially, the harmonic bond network connecting these particles is statically quenched at initialization. This represents the membrane as a 2D elastic solid rather than a purely fluid bilayer with lateral diffusion. This deliberate abstraction is justified by the disparate timescales of our system; the nanosecond duration of our simulated mechanical shock is orders of magnitude faster than lateral lipid diffusion (~ 1 *µ*m^2^*/*s). Consequently, the membrane topology is effectively frozen during rupture, allowing us to directly isolate the mechanical impact of quenched topological disorder (Seeded vs. Spontaneous). It is critical to note that under biological conditions, true lateral diffusion would permit dynamic domain reorganization. Under applied strain, nanopore nucleation preferentially initiates at the boundary interfaces between the rigid domains and the fluid matrix due to mechanical mismatch. If fluid mobility were not quenched, the sterol-rich domains could dynamically rearrange and flow to dissipate this local stress, effectively blunting crack tips at these interfaces. Therefore, the yielding mechanism observed in our quenched elastic solid represents a worst-case scenario, suggesting that the 39% topological reinforcement reported here is a highly conservative lower bound for the toughness of a fully fluid composite PDEV. We acknowledge that at this small scale, extreme curvature constraints artificially influence the absolute thermodynamic barrier to phase separation and the baseline bending rigidity ^21^.

Consequently, the critical rupture tensions reported here (e.g., 367.0 ± 0.7 mN m^−1^) must be interpreted as theoretical comparative indices rather than absolute physiological values. Furthermore, the linear constitutive mapping (Equation 1) used to derive these tensions is a first-order phenomenological approximation. The physical condensing effect of sterols is highly non-linear and typically plateaus at specific mole fractions. Our model assumes that the genetic parameters *G*_*SMT*_ and *G*_*FAD*_ operate strictly within a sub-saturating regime where these non-linear cooperativities remain negligible. For context, macroscopic micropipette aspiration experiments typically report lysis tensions of 5 to 15 mN m^−1^ for fluid phosphocholine bilayers, and up to 30 mN m^−1^ for sterol-rich raft mixtures ^12^. The order-of-magnitude discrepancy between these experimental baselines and our theoretical values is a well-characterised artifact of the computational timescale. Rupture is a kinetically driven process; the nanosecond loading rates inherent to molecular dynamics simulations afford insufficient time for thermal pore nucleation, inevitably yielding much higher rupture thresholds than the quasi-static loading of macroscopic experiments ^27^. However, classical kinetic theories of membrane failure dictate that critical rupture tension scales logarithmically with the applied loading rate (*γ*_*c*_ ∝ ln(*r* _*f*_)). Because this logarithmic scaling law applies universally to the underlying lipid continuum, a reduction to quasi-static experimental speeds shifts the absolute tensions downward while preserving the relative structural hierarchy. Consequently, the 39% mechanical reinforcement observed between the Ripening and Defence states remains a mathematically robust comparative index, independent of this kinetic bias. While conducting simulations across a broad spectrum of loading rates to extrapolate a quasi-static yield limit via Arrhenius kinetics would yield a more absolute metric of toughness, such an extensive parameter sweep across multiple topologies is computationally prohibitive for a model of this scale. Because the kinetic bias applies uniformly across all tested phenotypes, our comparative 39% reinforcement hypothesis remains robust.

Secondly, we simplified the complex plant lipidome into representative proxies (POPC/Phytosterol/PLD-products). While our use of POPC as the fluid matrix is biophysically justified by its asymmetric saturation ^22^, native plant membranes contain a rich diversity of minor components, notably glycosyl inositol phosphoryl-ceramides (GIPCs) and native membrane proteins ^23^. The exclusion of these highly polar, structurally asymmetric sphingolipids means our model inherently limits the physiological accuracy of domain condensation. *In vivo*, the bulky glycosylated headgroups and saturated acyl chains of GIPCs act as profound domain stabilisers and line-actants. Theoretically, the inclusion of GIPCs would necessitate an expansion of our constitutive mapping (Equation 1) to include a robust stiffening term (e.g., +*γG*_*GIPC*_). The dense hydrogen-bonding network provided by their large glycosylated headgroups would substantially elevate the baseline area compressibility modulus and further deepen the thermodynamic well of liquid-ordered domain separation.

In a coarse-grained framework, their inclusion would necessitate significantly deeper Lennard-Jones interaction energies (*ε*_*Lo*_) and would drastically increase the line tension at the liquid-ordered/liquid-disordered (*L*_*o*_*/L*_*d*_) interface. In a theoretical sensitivity analysis, artificially elevating the Lennard-Jones interaction energy (*ε*_*Lo*_) to approximate the massive line tension of GIPCs would aggressively drive domain coalescence. This heightened energetic penalty at the liquid-ordered/liquid-disordered (*L*_*o*_/*L*_*d*_) interface would restrict the entropic dispersal of domains, fundamentally shifting the percolation threshold. Under these parameters, the plant cell could achieve the monolithic limit (*S*_*max*_ ≈ 50%) at much lower bulk sterol/sphingolipid concentrations, while the Seeded composite domains would exhibit vastly enhanced resistance to shear. Therefore, the composite rigid domain formation we observe using our POPC proxy likely represents a highly conservative baseline estimate of the true mechanical reinforcement present in native PDEVs.

Future experimental validation is critical to translating these theoretical indices into physiological parameters. Specifically, we propose that atomic force microscopy (AFM) nano-indentation assays could be performed on PDEVs isolated from a wild-type plant model (Ripening baseline) versus those isolated from the same species subjected to controlled abiotic stress (Defence state). Quantifying the relative difference in vesicle bending moduli and yield forces between these two populations *in vitro* would definitively test the 39% reinforcement hypothesis predicted by our multiscale model.

Despite these abstractions, the engineering implications are clear. Current synthetic liposome strategies often focus on maximising bilayer rigidity, leading to stability-permeability trade-offs. Our findings offer an alternative, bio-inspired blueprint: the design of *composite* vesicles. By engineering delivery vectors that mimic the phase-separated topology of the Defence state, nanomedicine could replicate the mechanical homeostasis of PDEVs, creating carriers that are tough enough to survive the stomach yet compliant enough to fuse with target cells in the gut.

## A EV model

### A.1 SCG-MD potentials and force fields

The supra-molecular coarse-grained (SCG) model assumes a particle density of *ρ* = 0.5 nm^−2^. The total potential energy *U*_*total*_ is defined as the sum of three terms:

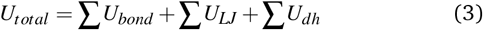

#### Elastic network

The membrane continuum is maintained via a harmonic bond network connecting nearest neighbours (cutoff *r*_*cut*_ = 2.5 nm). The spring constant *k*_*bond*_ is derived locally from the mapped area compressibility modulus *K*_*A*_(**G**):

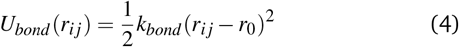

where *k*_*bond*_ ≈ 0.5*K*_*A* 31_. For the Defence state (*K*_*A*_ ≈ 360 mN m^−1^), this results in a stiffer lattice compared to the Ripening state (*K*_*A*_ ≈ 200 mN m^−1^).

#### Phase separation (Lennard-Jones)

To drive phase separation, we differentiate interactions between lipid species (*S*_*sat*_, *S*_*unsat*_). Sterol-rich domains are modelled using a truncated Lennard-Jones potential with a depth *ε* tuned to the condensing effect:

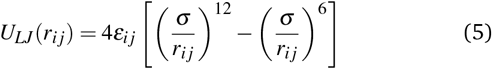

where *ε*_*sterol* −*sterol*_ = 15.0 kJ mol^−1^ (≈6*k*_*B*_*T*) to represent the deep thermodynamic well of raft formation at *T* = 300 K, while *ε*_*mix*_ = 2.5 kJ mol^−1^ for fluid interactions. It must be noted that because the bond network is statically quenched, these Lennard-Jones potentials do not drive dynamic phase separation during the pulling phase, but rather define the cohesive strength of the pre-initialized domains.

#### Electrostatics (Debye-Hückel)

pH-dependent interactions are modelled using a screened Coulomb potential:

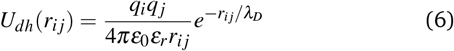

where the Debye length *λ*_*D*_ is set to 0.8 nm. This screening length was chosen to approximate the high ionic strength of the human gastrointestinal tract (≈ 150 mM), providing a realistic electrostatic environment for the simulated acid shock. The charge *q*_*i*_ of phosphatidic acid (PA) particles is modulated dynamically based on the environmental pH relative to the lipid pKa (approx 3.5 for PA).

### A.2 Topological initialisation algorithms

The difference between the two topological modes lies in the initial assignment of particle identities:

#### 1. Spontaneous (Entropic)

Particle identities (e.g., `Sterol, POPC`) are assigned via a uniform random distribution function ‘numpy.random.shuffle’. This simulates a membrane formed by pure entropic mixing without active cellular sorting mechanisms.

#### 2. Seeded (Active sorting)

To simulate Golgi-derived sorting, we utilise a *cluster growth algorithm*:

1. *N*_*seeds*_ nucleation sites are selected randomly on the spherical mesh.
2. A breadth-first search (BFS) traverses the neighbour list from each seed.
3. Neighbours are assigned the Sterol identity until the target fraction *φ*_*sterol*_ is reached.
4. Remaining particles constitute the fluid matrix.

This ensures that the Defence state begins with consolidated liquid-ordered (*L*_*o*_) domains, mimicking the effect of lipid flippases and raft-associated proteins.

### A.3 Quantification of morphology

#### Percolation threshold (*S*_*max*_)

We quantify the connectivity of the rigid phase using a Union-Find algorithm on the sterol subgraph, applying classical percolation theory to evaluate the lattice threshold ^32^. *S*_*max*_ is defined as the fraction of total membrane particles belonging to the largest connected rigid component. A transition from *S*_*max*_ ≈0.15 (Dispersed) to *S*_*max*_ ≥0.45 (Monolithic limit for a 50% rigid composition) indicates the formation of a global percolating network..

#### Spatial heterogeneity (Moran’s I)

To distinguish between random noise and ordered domains, we compute Moran’s *I* autocorrelation index, a standard measure of spatial clustering ^33^::

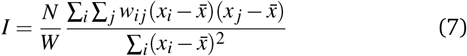

where *w*_*i j*_ is the spatial weight matrix (1 for neighbours, 0 otherwise). *I*≈ 0 indicates random mixing (Spontaneous), while *I*→ 1 indicates high spatial clustering (Seeded).

## B Simulation setup

### Integrator and thermodynamic

Temperature *T* = 300 K; Langevin friction *γ* = 1.0 ps^−1^; time step 4 fs.

### Geometry and resolution

Vesicle radius *R* = 20 nm; surface particle density *ρ* = 0.5 nm^−2^. Bond network constructed from neighbor pairs within 2.5 nm.

### Nonbonded interactions

Cutoff 3 nm; Lennard–Jones *ε* = 15.0 kJ mol^−1^ (*L*_*o*_) vs 1.0 kJ mol^−1^ (*L*_*d*_); screened Coulomb with Debye length 0.8–1.0 nm.

### External loading and failure

Pressure ramp parameter *k*_pressure_ incremented by 0.5 kJ mol^−1^nm^−1^ per 500 steps; failure criterion at 10–15% radial expansion (irreversible plastic regime).

### Initialisation modes and randomness

Seeded (cluster-growth) and Spont (random assignment) initialisations are used; distinct random seeds and initial domain assignments across replicates.

### Code availability

OpenMM implementations and figure production scripts are available at https://github.com/Jagirhussan/Plant-derived-extracellular-vesicles.

## C Statistical tables

**Table 1.**
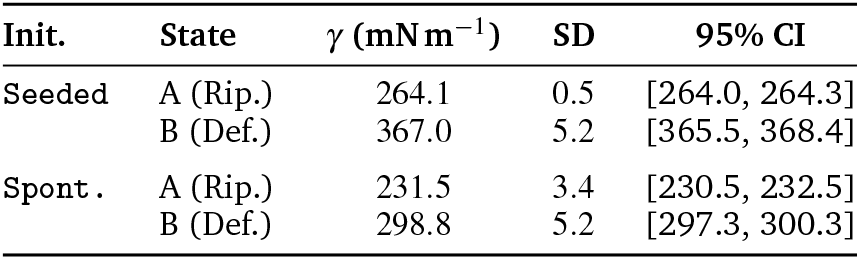
Rupture tension statistics (*n* = 50 per group; mean ± SD, 95% CI).

**Table 2.** Percolation fraction statistics (*n* = 50 per group; mean ± SD, 95% CI).

| Init. | State | Mean | SD | 95% CI |
| --- | --- | --- | --- | --- |
| Seeded | A (Rip.) | 0.019 | 0.008 | [0.017, 0.021] |
|  | B (Def.) | 0.377 | 0.084 | [0.353, 0.401] |
| Spont. | A (Rip.) | 0.005 | 0.001 | [0.005, 0.006] |
|  | B (Def.) | 0.495 | 0.004 | [0.494, 0.496] |

**Table 3.** Moran’s *I* statistics (*n* = 50 per group; mean ± SD, 95% CI).

| Init. | State | $I$ | SD | 95% CI |
| --- | --- | --- | --- | --- |
| Seeded | A (Rip.) | 0.610 | 0.026 | [0.603, 0.618] |
|  | B (Def.) | 0.610 | 0.021 | [0.604, 0.616] |
| Spont. | A (Rip.) | 0.003 | 0.011 | [-0.000, 0.006] |
|  | B (Def.) | 0.001 | 0.008 | [-0.002, 0.003] |

**Table 4.** Welch two-sample *t*-tests (Defence vs Ripening) by initialisation mode.

| Mode | $\Delta$<br>(mN m<sup>-1</sup>) | $t$ | $p$ | Cohen's $d$ |
| --- | --- | --- | --- | --- |
| Seeded | 102.8 | 140.64 | $2.20 \times 10^{-66}$ | 28.13 |
| Spont. | 67.3 | 76.20 | $5.41 \times 10^{-80}$ | 15.24 |

## Notes

https://github.com/Jagirhussan/Plant-derived-extracellular-vesicles

### Competing Interest Statement

The authors have declared no competing interest.

### Funding

- Ministry of Business, Innovation and Employment: UOA2510

